# L3R-seq: A long-read 3’RACE approach for deep quantitative analysis of RNA processing

**DOI:** 10.64898/2026.05.20.726719

**Authors:** Akihito Mamiya, Mizuki Takenaka, Munetaka Sugiyama

**Affiliations:** Department of Botany, Graduate School of Science, Kyoto University; Department of Biology, Graduate School of Science, Kobe University, Japan; Department of Biological Sciences, Graduate School of Science, The University of Tokyo, Japan

**Keywords:** Long-read, ONT, PacBio, UMI, RNA editing, polyadenylation, plant mitochondria

## Abstract

Long-read sequencing technologies offer the potential to capture multiple RNA processing events within single molecules, but standard protocols suffer from quantification biases and sequencing errors that limit their utility for precise analysis. Here, we describe **Long-read 3’ RACE-seq** (**L3R-seq**), a targeted long-read sequencing method that ligates a unique molecular identifier (UMI)-containing adapter to the 3’ end of RNA molecules prior to reverse transcription and PCR amplification. By grouping cDNA reads sharing the same UMI and generating a consensus sequence for each original RNA molecule, L3R-seq corrects random sequencing errors and mitigates PCR-duplicate-driven quantification biases. The method enables simultaneous, per-molecule analysis of RNA editing, 3’ end cleavage and trimming, and polyadenylation status. Along with a step-by-step protocol for library preparation and sequencing with the Oxford Nanopore Technologies (ONT) platform, we describe an accompanying bioinformatic pipeline for consensus generation and extraction of RNA features. As an example, we apply L3R-seq to the mitochondrial mRNA *ccmC* from *Arabidopsis thaliana*, a transcript subject to extensive C-to-U editing and non-canonical 3’ end processing. The workflow is readily adaptable to other RNAs targets and is transferable to the Pacific Biosciences (PacBio) platform.

## 1 Introduction

RNA maturation requires multiple processing steps, including editing, splicing, base modification, terminal cleavage, trimming, and polyadenylation. These processes often influence one another, adding to the complexity of RNA metabolism *in vivo*. Next-generation short-read sequencing has been widely adopted to investigate these RNA processing events ***[1]***. However, due to constraints in sequencing read length, these approaches are mostly incapable (with a few exceptions ***[2–4]***) of directly recording the long-range associations of multiple processing events. Recently, high throughput long-read sequencing technologies from Oxford Nanopore Technologies (ONT) and Pacific Biosciences (PacBio) have emerged ***[1, 5–7]***, making it possible to address this limitation. Between the two platforms, ONT ***[6]*** has the advantage of (i) accommodating an exceptionally wide range of read lengths, (ii) flexible experimental scale, including small-scale pilot studies, and (iii) direct detection of base modifications in both DNA and RNA; however, (iv) its raw-read accuracy remains suboptimal for precise detection and quantification of single-nucleotide changes (typically ∼Q20 for simplex DNA reads). PacBio ***[7]***, by contrast, provides (i) high-fidelity (HiFi) reads with exceptional quality (Q30+); however, it (ii) accommodates a narrower range of read lengths than ONT, particularly under concatenation-based HiFi protocols (*see* **Note 1**), (iii) is optimized for large-scale experiments, and (iv) offers fewer capabilities for base-modification detection.

Despite their promise, standard long-read RNA sequencing protocols on both ONT and PacBio platforms can distort isoform and transcript-end representation ***[8]***, complicating accurate quantification of transcript termini, RNA editing, and other processing events. These distortions arise from biases at multiple stages of library preparation, including oligo(dT) mispriming at internal adenine stretches, preferential PCR amplification of shorter fragments, and size-dependent losses during bead-based DNA purification steps. Platform-specific artifacts further compound these issues. In ONT sequencing, shorter molecules translocate through pores more rapidly, leading to their preferential sequencing at the expense of longer fragments. Random sequencing errors further limit precise detection of single-nucleotide changes. PacBio HiFi sequencing largely circumvents read accuracy limitations, though its concatenation-based protocols impose their own read-length constraints (*see* **Note 1**).

To address these challenges, we introduce <u>L</u>ong-read <u>3</u>’ <u>R</u>ACE-seq (**L3R-seq**), a long-read DNA sequencing method for analyzing 3’ RACE amplicons (**Fig. 1**). This method ligates the 3’ end of RNA molecules to an adapter originally developed by Scheer et al. ***[9]***, which contains a 15 nucleotide random region, called the unique molecular identifier (UMI) ***[10]***. The UMI enables grouping of cDNA reads derived from the same RNA molecule. Generating a consensus among them leads to the correction of random sequencing errors and mitigates PCR-duplicate-driven biases in quantification. Although the same adapter has been used to study 3’ RNA extremities with Illumina sequencing ***[9]*** and with ONT ***[11]***, methods that utilize its UMI in improving read accuracy have not been developed. Here, we outline the experimental and bioinformatic techniques (**Fig. 2**) to obtain the consensus sequence, map to a reference, and call RNA features on a per-molecule basis (i.e. editing, 3’ terminal position and 3’ extensions).

**Fig. 1.**
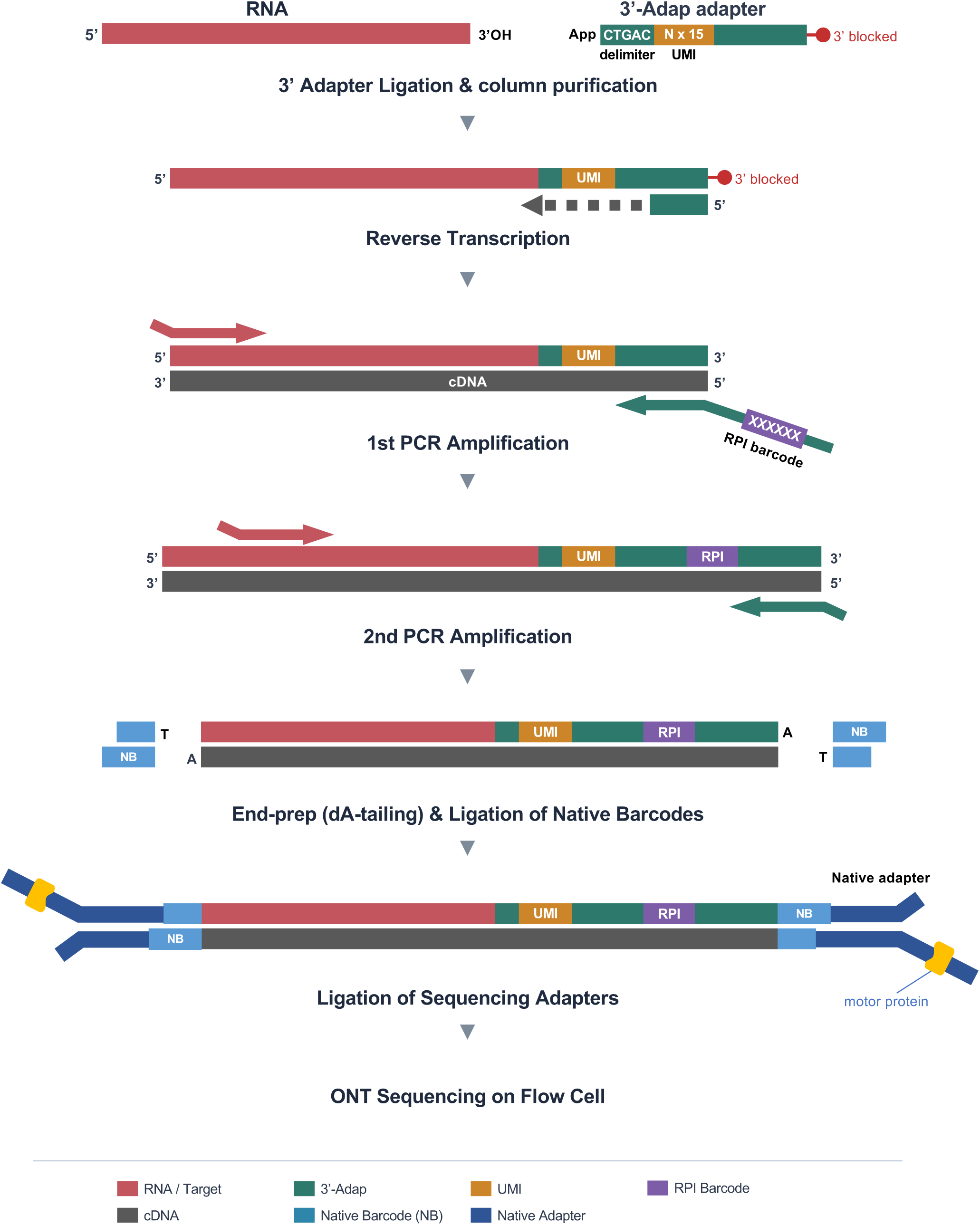
L3R-seq library preparation. Total RNA is ligated to the 3’-Adap adapter containing a unique molecular identifier (UMI). After column purification, cDNA is synthesized using SMARTScribe reverse transcriptase. Target 3’ regions are amplified by nested PCR with RPI barcode primers. The resulting amplicons undergo ONT library preparation using the Native Barcoding Kit V14 (e.g. SQK-NBD114.24): end-prep adds dA-tails, native barcodes (NB) are ligated via T-overhang complementarity, and sequencing adapters are ligated to both ends. Libraries are sequenced on a nanopore flow cell.

**Fig. 2.**
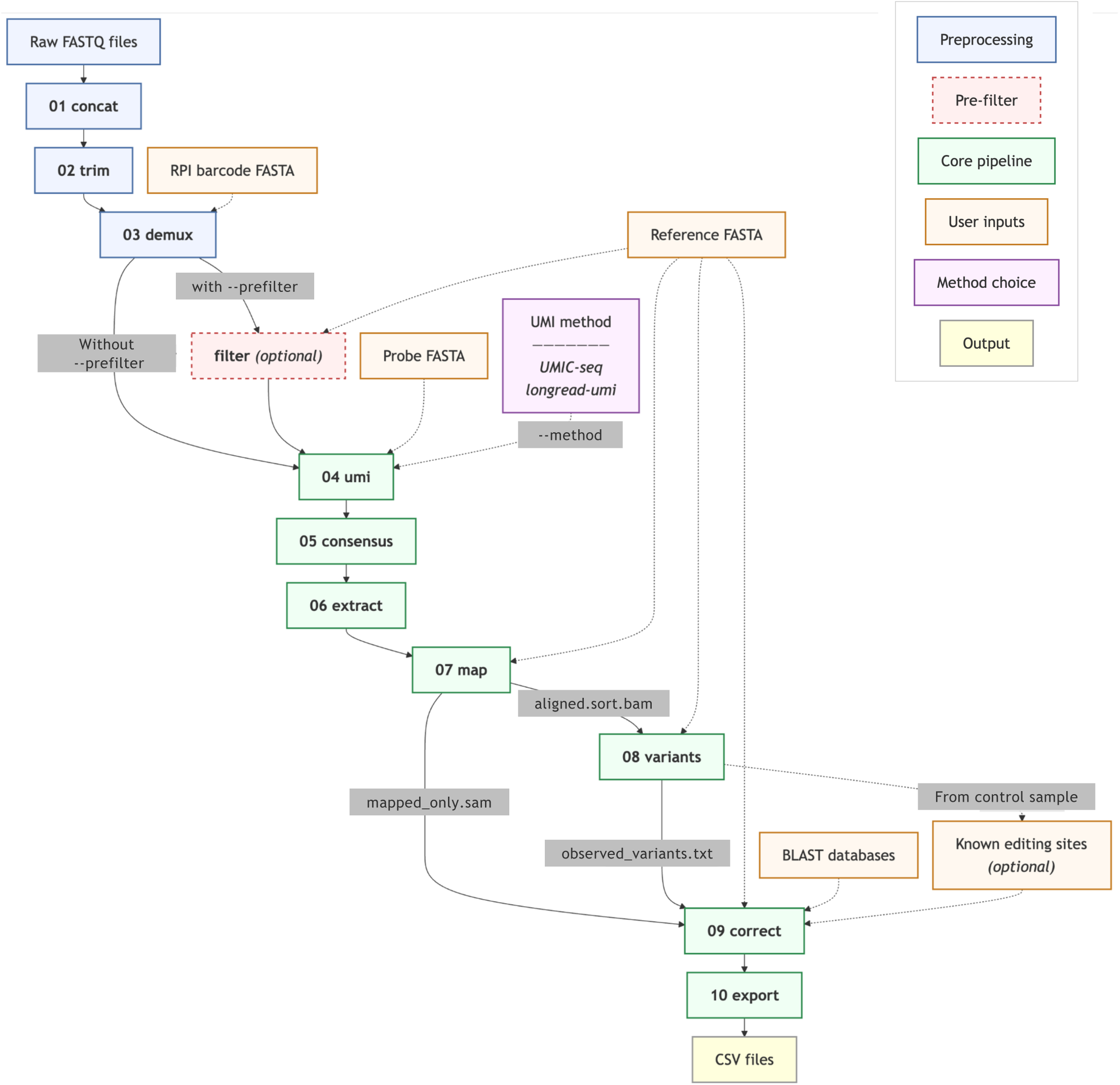
The L3Rseq bioinformatic pipeline. Numbered boxes represent L3Rseq subcommands. Raw FASTQ reads are concatenated (01 concat), adapter-trimmed (02 trim), and demultiplexed by RPI barcodes (03 demux). An optional pre-filtering step retains only on-target reads (filter). UMIs are extracted and reads are clustered by hierarchical clustering using UMIC-seq or longread-umi (04 umi), and per-cluster consensus sequences are generated by iterative Racon polishing (05 consensus). The target region is extracted by trimming flanking primer and adapter sequences (06 extract), and consensus sequences are mapped to the reference with minimap2 (07 map). Single-nucleotide variants are called with LoFreq to identify C-to-U RNA editing sites (08 variants). 3’ tail alignments are corrected by tolerating C-to-T mismatches at known editing positions, with optional BLAST searches to detect translocations (09 correct). Per-molecule annotations are exported to CSV files (10 export). Rounded boxes indicate external inputs; the dashed box indicates an optional step.

This protocol is especially advantageous when oligo(dT)-primed reverse transcription is not appropriate for cDNA synthesis—for instance, when RNAs of interest may lack long poly(A) tails, or when the status of RNA 3′ termini itself is the focus of the study. As an example, we analyzed the mitochondrial mRNA *ccmC* from the model plant *Arabidopsis thaliana* (**Fig. 3**). mRNAs transcribed from the plant organellar genomes (mitochondrion and chloroplast) are known to undergo C-to-U base editing mediated by pentatricopeptide repeat (PPR) proteins ***[12]***, and *ccmC* is one of the most densely edited with 28 editing sites ***[13]***. In addition, 3’ poly(A) tails on plant organellar mRNAs, including *ccmC*, are typically short and are rarely detected, likely reflecting the organelles’ inheritance of bacterial-type gene expression systems ***[14]***. Moreover, the *ccmC* mRNA lacks a stop codon, as a consequence of endonucleolytic cleavage within the ORF, followed by exonucleolytic trimming of the 3’ end ***[15]***. Altogether these properties make *ccmC* a compelling target to simultaneously examine the multiple processing events (i.e. RNA editing, 3’ end cleavage/trimming, and polyadenylation) within each single RNA molecule. Sequencing was performed on the ONT platform, but the workflow is readily transferable to PacBio (*see* **Note 1**).

**Fig. 3.**
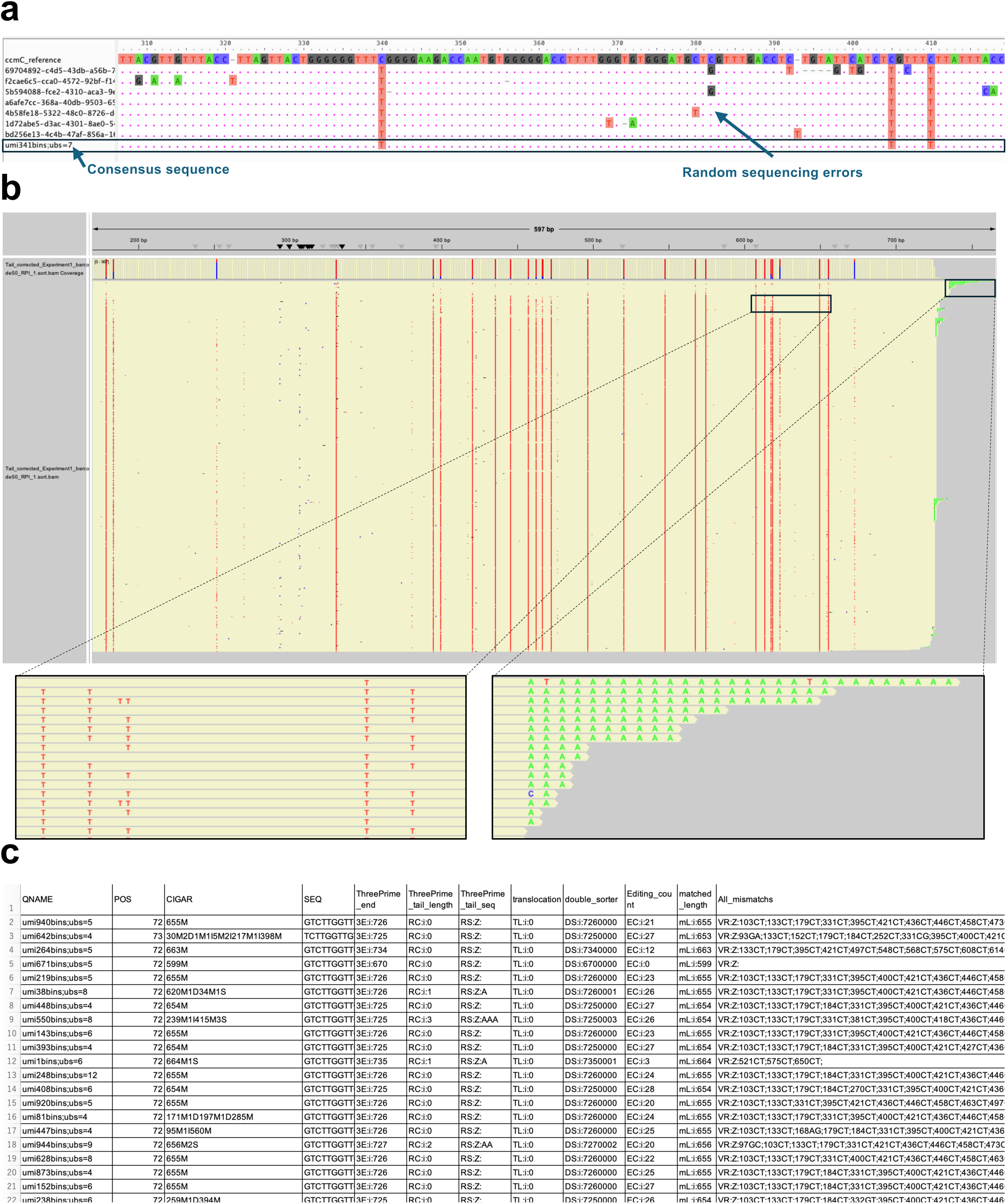
Representative L3R-seq output for *Arabidopsis* mitochondrial *ccmC* mRNA. **(a)** Raw nanopore reads within a single UMI cluster (umi341bins; 7 reads) were aligned to the *ccmC* genomic reference using the AliView software ***[32]***. Colored bases indicate positions differing from the reference, including C-to-U RNA editing sites. These reads share the same UMI, deriving from a single original RNA molecule, and are collapsed into one high-accuracy consensus sequence (section 3.7, step 3), as shown at the bottom. **(b)** An IGV view of the tail-corrected consensus sequences mapped to the *ccmC* reference. Each horizontal bar represents one consensus molecule. Positions colored in red indicate C-to-U editing events relative to the genomic reference; non-templated 3′ poly(A) tails are visible in green. Coverage tracks are shown above. Expanded insets below highlight two key features: (left) a close-up of representative C-to-U editing sites, where T bases in the consensus reads mark positions at which the transcript has been edited from C to U relative to the genomic sequence; (right) the 3′ terminal region, showing non-templated tail extensions predominantly composed of A residues, consistent with short poly(A) additions characteristic of plant mitochondrial mRNAs. **(c)** Select rows are shown from the per-molecule annotation table generated by L3Rseq export (section 3.9, step 3). Each row represents one original RNA molecule. Columns include alignment information (QNAME, POS, CIGAR, SEQ), 3′ end position on the reference (ThreePrime_end), 3′ tail length and sequence (ThreePrime_tail_length, ThreePrime_tail_seq), translocation status, C-to-T editing count, matched alignment length, and all detected single-nucleotide variants (All_mismatches).

Several established long-read workflows also leverage UMIs to build per-molecule consensus sequences. The longread_umi ***[16]*** pipeline and its derivatives (*e.g*. ***[17]***) implement dual UMI tagging by introducing UMI-containing forward and reverse primers at the first two cycles of PCR. R2C2 + UMI ***[18]*** combines this strategy with R2C2 ***[19]***, a technique in which circularized double-stranded cDNAs undergo rolling-circle amplification to generate tandem repeats that could be collapsed into an ultra-accurate consensus. In contrast, the L3R-seq method ligates a single UMI adapter to the 3’ end of RNA. This enables precise mapping of the 3’ end, avoids amplification failures associated with long, tag-bearing PCR primers, permits the use of multiple gene-specific forward primers to increase the specificity of the PCR product, and simplifies the workflow by eliminating cDNA circularization and rolling-circle amplification. Given the recent improvements in ONT read accuracy ***[5, 6]***, repeat-based consensus strategies may be unnecessary unless exceptionally high per-base accuracy is required. Limitations of L3R-seq include the inability to recover full-length reads spanning the 5′ end and, because it is cDNA-based, the lack of direct RNA-modification detection. Overall, L3R-seq offers a simple workflow for interrogating long-range associations among RNA-processing events within single RNA molecules.

## 2 Materials

### 2.1 3’ Adapter ligation and clean up

1. RNA extracted by standard methods (e.g. TRIzol)
2. 10 U/µL T4 RNA Ligase 1 (ssRNA Ligase) (NEB, M0204)
3. 10x T4 RNA Ligase Reaction Buffer (Supplied with NEB, T4 RNA Ligase 1)
4. 50% PEG 8000 (Supplied with NEB, T4 RNA Ligase 1)
5. DMSO (Molecular biology grade)
6. 3’-Adap oligonucleotide: /5rApp/CTGACNNNNNNNNNNNNNNNTGGAATTCTCGGGTGCCAAGGC/3ddC/ (Integrated DNA Technologies)
7. NucleoSpin RNA Clean-up XS (MACHEREY-NAGEL)
8. 1.5 mL DNA LoBind tubes (Eppendorf)
9. Incubator for 1.5 mL tubes at 16°C and 65°C
10. Nuclease-free water

### 2.2 cDNA synthesis

1. 100 U/µL SMARTScribe Reverse Transcriptase (Clontech)
2. 5x First-strand Buffer (Supplied with Clontech, SMARTScribe RTase)
3. 20 mM DTT (Supplied with Clontech, SMARTScribe RTase)
4. dNTP Mix (10 mM each)
5. 3’-RT primer: GCCTTGGCACCCGAGAA (Integrated DNA Technologies)
6. PCR tubes
7. PCR thermal cycler
8. Nuclease-free water

### 2.3 PCR amplification and gel electrophoresis

1. PrimeSTAR Max DNA Polymerase (Takara)
2. PCR primers for 1st PCR

i. Gene-specific forward primer for 1st PCR
ii. Adapter-specific reverse primer (e.g. Illumina RPI primer: CAAGCAGAAGACGGCATACGAGAT**XXXXXX**GTGACTGGAG TTCCTTGGCACCCGAGAATTCCA) ***[9]***
3. PCR primers for 2nd PCR

i. Gene-specific forward primer for 2nd PCR
ii. Adapter-specific reverse primer (e.g. 3’ Adapter_Rv_2: CAAGCAGAAGACGGCATACGAGAT)
4. PCR tubes
5. PCR thermal cycler
6. Exonuclease I (NEB, M0293)
7. Standard equipment for agarose gel electrophoresis (Agarose, TAE buffer, electrophoresis power supply, casting tray, well combs and gel box)
8. EzFluoroStain DNA (ATTO)
9. SPRIselect beads (Agencourt) or a PCR clean-up column kit (Wizard SV Gel and PCR Clean-Up System, Promega)
10. Qubit 4 Fluorometer (ThermoFisher Scientific)
11. Qubit dsDNA BR Assay Kit (Invitrogen)

### 2.4 ONT library preparation

1. Native Barcoding Kit V14 (SQK-NBD114, ONT) (*see* **Note 2**)
2. NEBNext Ultra II End repair/dA-tailing Module (NEB, E7546)
3. NEB Blunt/TA Ligase Master Mix (NEB, M0367)
4. NEBNext Quick Ligation Module (NEB, E6056)
5. SPRIselect (or AMPure XP) magnetic beads (Agencourt) (*see* **Note 3**)
6. 99.5% Ethanol
7. Magnetic stand for bead purification of DNA
8. Hula mixer (or rotator mixer) for DNA purification with magnetic beads
9. Incubator at 37°C
10. ONT flow cell (e.g. R10.4.1 MinION flow cell)
11. ONT sequencer (e.g. MinION Mk1D)
12. 50 mg/ml Bovine Serum Albumin (BSA) (Invitrogen UltraPure BSA 50 mg/ml, AM2616)
13. Qubit 4 Fluorometer (ThermoFisher Scientific)
14. Qubit dsDNA HS Assay Kit (Invitrogen)

### 2.5 Hardware requirements

Basecalling with Dorado benefits substantially from an NVIDIA GPU. The L3R-seq analysis pipeline described below is CPU-only and does not require a GPU.

We developed and tested L3R-seq on the following systems:

1. Intel Core i7-11700 CPU, NVIDIA GeForce RTX 3060 12GB GPU, 128GB RAM.
2. AMD Ryzen 9 9950X CPU, NVIDIA GeForce RTX 5090 32GB GPU, 128GB RAM.

### 2.6 Software requirements

1. MinKNOW software (ONT)
2. (*optional*) EPI2ME software (ONT)
3. L3Rseq, a software suite of command-line tools developed to accompany the L3R-seq sequencing method, is publicly available as a pre-built Docker image through the GitHub Container Registry (ghcr.io/akihito-mamiya-del/l3rseq:latest), enabling straightforward deployment across computing environments. The software and documentation can be found at https://github.com/akihitomamiya-del/L3R-seq.
4. A Debian-based Linux environment. L3Rseq has been tested on native Ubuntu 24.04, an Ubuntu 24.04-based Docker container on Windows 10 with WSL, and a Debian-based Node.js 20 container. Other Debian-based distributions are expected to be compatible.

## 3 Methods

### 3.1 3’ Adapter Ligation and RNA clean-up

1. A 5’-riboadenylated DNA oligonucleotide (3’-Adap) is ligated to the 3’ ends of RNA ***[9, 11]***.

a. In a 1.5 mL DNA LoBind tube, take 2 µg of total RNA and add nuclease-free water to a final volume of 6 µL.
b. Add 1 µL of 10 µM 3’-Adap.
c. Denature sample for 3 min at 65°C and immediately cool on ice for 3 min.
d. Place the tube at room temperature and add the following and mix by pipetting.

i. 2 µL 10x T4 RNA Ligase Reaction Buffer
ii. 2 µL DMSO
iii. 1 µL T4 RNA ligase
iv. 8 µL 50% PEG 8000
e. Incubate at 16°C, overnight.
2. Perform RNA clean up with a column kit (NucleoSpin RNA Clean-up XS) following the manufacturer’s protocol. Elute in 10 µL Nuclease-free water.

### 3.2 cDNA Synthesis

1. Synthesize cDNA with a primer complementary to 3’-Adap.

a. In a 0.2 mL PCR tube, take 4 µL of RNA from the previous step (the remaining eluate may be stored at –80°C for replicate or additional reactions).
b. Add 1 µL of 20 µM 3’-RT primer.
c. Denature sample for 3 min at 72°C and immediately cool on ice for 3 min.
d. Add the following, mix and spin down.

i. 2 µL of 5x First-Strand Buffer
ii. 1 µL dNTP Mix (10 mM each)
iii. 1 µL 20 mM DTT
e. Add 1 µL of 100 U/µL SMARTScribe RTase, mix and spin down.
f. In a PCR thermal cycler, incubate at 42°C for 90 min, then terminate the reaction by heating at 70°C for 15 min.
2. (*optional*) Dilute the cDNA solution by adding 40 µL of nuclease-free water.

### 3.3 PCR

1. Perform the 1st PCR with a gene-specific forward primer and an adapter-specific reverse primer containing sample-specific barcodes.

a. In a PCR tube, add the following and mix.

i. 1 µL cDNA
ii. 1 µL 5 µM Forward primer
iii. 1 µL 5 µM Reverse primer
iv. 9.5 µL Nuclease-free water
v. 12.5 µL PrimeSTAR Max Premix (2x)
b. In a PCR thermal cycler, run the following program.

i. 98°C, 2 min
ii. {98°C, 10 sec; 55°C, 5 sec; 72°C, 2 min} × 15 cycles
iii. 72°C, 4 min, cool down
2. Exonuclease digestion of the PCR primers. Residual primers from the 1st PCR are digested to prevent carryover into the nested 2nd PCR and reduce non-specific amplification.

a. Dilute Exonuclease I in PrimeSTAR Max Premix with nuclease-free water to a final concentration of 1U/µL Exonuclease I, 1x PrimeSTAR Max Premix.
b. Add 4 µL 1U/µL Exo I (in 1x PrimeSTAR Max premix) to 20 µL of the 1st PCR reaction.
c. Incubate at 37°C for 1h.
d. Stop the reaction by incubating at 80°C for 20 min.
3. Perform the 2nd PCR with a second gene-specific forward primer and an adapter-specific reverse primer.

a. In a PCR tube, add the following and mix.

i. 5 µL of Exonuclease I-treated 1st PCR reaction
ii. 1 µL Forward primer
iii. 1 µL Reverse primer
iv. 5.5 µL Nuclease-free water
v. 12.5 µL PrimeSTAR Max Premix (2x)
b. In a PCR thermal cycler, run the following program.

i. 98°C, 2 min
ii. {98°C, 10 sec; 55°C, 5 sec; 72°C, 2 min} × 20 cycles
iii. 72°C, 4 min, cool down
4. Run agarose gel (typically 1%) electrophoresis. The use of highly sensitive dyes (e.g. ATTO EzFluoroStain DNA) is recommended for visualization of the full range of PCR products. Typically, a primer dimer band will appear ∼ 100bp.
5. Size select against the PCR dimers and clean up the PCR products with gel extraction using a column kit (e.g. Wizard SV Gel and PCR Clean-Up System (Promega)). Alternatively, magnetic beads (SPRIselect) can be used. The concentration of beads (typically 0.5 ∼ 0.8x) should be determined depending on the desired size threshold ***[20]***. Amplicons that will be assigned the same native barcodes in the next library preparation step could be pooled together at this stage.
6. Quantify the purified amplicons with the Qubit fluorometer. 200 fmol (130 ng for 1 kb amplicons) in 12.5 µL is required for each native-barcoded sample in library preparation.

### 3.4 ONT Library Preparation

For ligation sequencing amplicons with the Native Barcoding Kit, download the latest protocol from the ONT [21] and follow the instructions; a brief version is provided below (*see* **Note 4**).

1. End-prep and bead purification

a. In a 1.5 mL DNA LoBind tube, mix the following.

i. 12.5 µL amplicon (∼ 200 fmol, *see **Note*** **5**)
ii. 1.75 µL Ultra II End-prep Reaction Buffer
iii. 0.75 µL Ultra II End-prep Enzyme Mix
b. Incubate at 20°C for 5 min.
c. Stop the reaction by incubating at 65°C for 5 min.
d. (Optional) Clean-up end-prepped samples with AMPure XP (or SPRIselect) beads, with two 80% Ethanol washes (*see **Note*** **5**).
2. Native barcode ligation

a. In a new DNA LoBind tube, mix the following (see ***Note* 6**).

i. 3 µL nuclease-free water
ii. 0.75 µL end-prepped amplicon
iii. 1.25 µL native barcode
iv. 5 µL Blunt/TA master mix
b. Incubate at room temperature for 20 min.
c. Stop the reaction by adding 2 µL 250 mM EDTA.
d. Pool the samples into one tube.
e. Purify the native-barcoded samples with 1x AMPure XP (or SPRIselect) beads with two 80% Ethanol washes and elute to 35 µL in nuclease-free water.
f. Quantify with Qubit.
g. Store the sample at 4°C overnight, if necessary.
3. Adapter ligation

a. In a new DNA LoBind tube, mix the following.

i. 30 µL pooled native-barcoded samples
ii. 5 µL native adapter
iii. 10 µL NEBNext Quick Ligation Reaction Buffer (5X)
iv. 5 µL Quick T4 DNA Ligase
b. Incubate at room temperature for 20 min.
c. Clean up the adapter-ligated sample with 0.5x AMPure XP (or SPRIselect) beads with two Short Fragment Buffer (SFB) washes (see **Note 7**) and elute to 15 µL Elution Buffer (EB) at 37°C.
d. Quantify with Qubit.

### 3.5 ONT sequencing

To sequence the ONT library, follow the instructions in the manuals for the Native Barcoding Kit ***[21]***, the sequencer and MinKNOW software; a brief version is provided below (see ***Notes* 4-5**).

1. Insert the flow cell to the sequencer, connect to PC and open the MinKNOW software.
2. Perform flow cell check.
3. Prime the flow cell.

a. Prepare priming mix (Flow Cell Flush (FCF) 1170 µL + Flow Cell Tether (FCT) 30 µL + 5 µL 50 mg/ml Bovine Serum Albumin (BSA)).
b. Load the priming mix to the flow cell.
4. Load the sequencing library.

a. Mix the following.

i. 37.5 µL Sequencing Buffer (SB)
ii. 25.5 µL Library Beads (LIB)
iii. 12 µL DNA library (typically adjusted to 100-200 fmol)
b. Load the library mix to the flow cell.
5. Start sequencing with the MinKNOW software. Set up live base calling if necessary (*see* **Note 8**).

### 3.6 Basecalling and Native Barcode Demultiplexing

Perform basecalling and native barcode demultiplexing with dorado, which is available within MinKNOW, EPI2ME, or as a standalone command-line tool (*see* **Note 9**). For standalone use, basecalling and barcode classification can be performed in a single step:

`dorado basecaller sup <POD5_DIR> --kit-name <e.g. SQK-NBD114-24> --min-qscore 10 > calls.bam`

The output is then demultiplexed into per-barcode BAM files:

`dorado demux --output-dir <OUTPUT_DIR> --no-classify calls.bam`

The `--no-classify` flag ensures that dorado uses the barcode assignments from the basecalling step rather than re-classifying the reads after trimming, which could interfere with barcode detection. Convert the per-barcode BAM files to FASTQ format for downstream processing with L3Rseq.

### 3.7 UMI clustering and consensus generation

The base-called, native-barcode-demultiplexed reads from section 3.6 are processed through the L3Rseq pipeline to generate a consensus sequence for each original RNA molecule. L3Rseq is invoked as a series of subcommands, each performing one stage of the analysis. Alternatively, the entire pipeline can be executed with a single command (`L3Rseq run`; *see* **Note 10**). All steps described below are run within the L3Rseq Docker container environment (section 2.6).

1. (Optional) Concatenate, trim, and demultiplex the reads (**Fig. 2** steps 01-03). If starting from raw MinKNOW output, three preprocessing steps are available. First, per-barcode fastq.gz files produced by MinKNOW are concatenated into a single file per ONT barcode (`L3Rseq concat`). Second, library preparation adapters are trimmed in three passes using cutadapt ***[22]*** (`L3Rseq trim`). Third, reads are demultiplexed by the RPI barcodes introduced during PCR (section 3.3) using cutadapt (`L3Rseq demux`). These steps use adapter and barcode sequences from the current protocol as defaults; users with different library preparation designs can override them with command-line flags (*see* **Note 11**). Skip these steps if using pre-demultiplexed data (e.g. by setting `--start-at 4` in `L3Rseq run`).
2. Extract UMIs and cluster reads (**Fig. 2** step 04). L3Rseq locates the 15-nt UMI within each read by aligning a probe sequence complementary to the adapter region flanking the UMI (3’-Adap). The UMI is extracted and reads sharing the same UMI—i.e. reads derived from the same original RNA molecule—are grouped by hierarchical clustering using UMIC-seq ***[23]***. A cluster test is first performed on a subset of reads to determine the optimal alignment score threshold, followed by full clustering of all reads at the selected threshold (*see* **Note 12**).

L3Rseq umi --input <DEMUX_DIR> --outdir <OUTPUT_DIR> --probe <probe.fasta>

1. Generate consensus sequences (**Fig. 2** step 05). Within each UMI cluster, reads are aligned with minimap2 ***[24]*** and polished through four iterative rounds of Racon ***[25]*** to produce a single high-accuracy consensus sequence per original RNA molecule (**Fig. 3a**). This step corrects random ONT sequencing errors and collapses PCR duplicates, yielding per-molecule sequences suitable for single-nucleotide variant detection (*see* **Note 13**).

L3Rseq consensus --input <INPUT_DIR> --outdir <OUTPUT_DIR>

### 3.8 Mapping to reference

The consensus sequences are trimmed to the target region and mapped to a reference sequence for downstream feature calling.

1. Extract the target region (**Fig. 2** step 06). Cutadapt ***[22]*** is used to trim the flanking primer and adapter sequences from each consensus, isolating the region of biological interest. For the *ccmC* analysis, the forward target sequence corresponds to the gene-specific primer and the reverse target sequence spans the adapter boundary, retaining the 3’ extension beyond the coding region. Users analyzing different targets should adjust these sequences with the `--target-fwd` and `--target-rev` flags (*see* **Note 14**). `L3Rseq extract --input <INPUT_DIR> --outdir <OUTPUT_DIR>`
2. Map to the reference (**Fig. 2** step 07). The extracted consensus sequences are aligned to the reference sequence using minimap2 with the `lr:hq` preset, appropriate for high-quality long reads such as polished consensus sequences. The output is converted to sorted BAM format and indexed with samtools ***[26, 27]***. After this step, each alignment represents a single original RNA molecule. Each alignment in the BAM output carries a CIGAR string, a field defined in the SAM format specification ***[26]*** that describes how the read aligns to the reference, encoding both the matched region and any 3’ sequence extending beyond the reference as soft-clipped bases (**Fig. 4**). `L3Rseq map --input <INPUT_DIR> --outdir <OUTPUT_DIR> --ref <reference.fasta>`

**Fig. 4.**
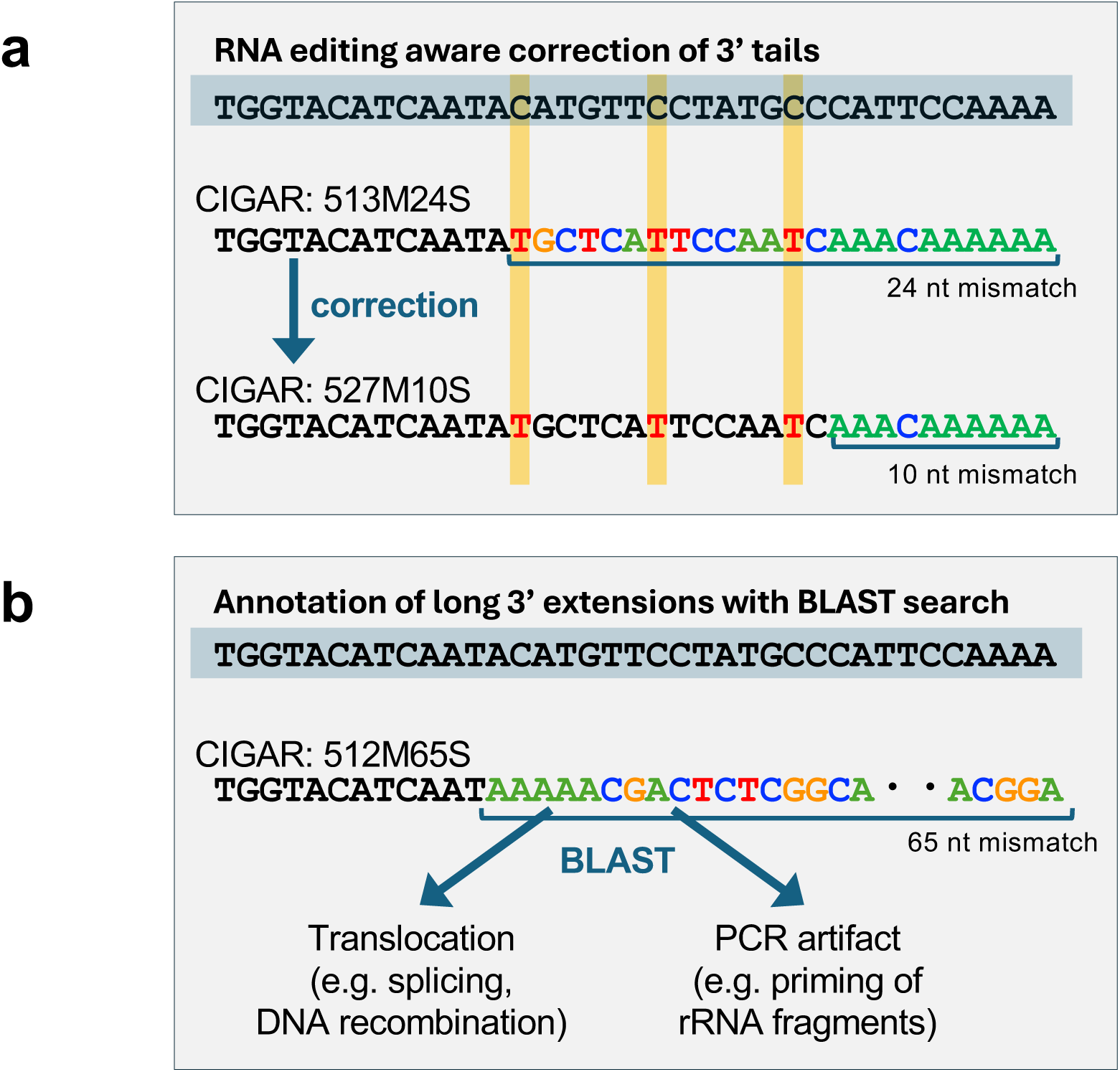
Schematic illustration of the 3′ tail correction and long 3′ extension annotation procedures in L3R-seq. **(a)** Correction of 3′ tails using known RNA editing patterns (section 3.9, step 2). The top row shows the genomic reference sequence near the 3′ end, with C-to-U RNA editing positions highlighted (yellow lines). Below, a consensus read is shown before and after tail correction. Before correction (CIGAR: 513M24S), the aligner soft-clips 24 nucleotides because C-to-U edited positions near the 3′ boundary appear as mismatches against the genomic reference. The walk correction algorithm tolerates C-to-T mismatches at known editing sites (yellow), extending the aligned region to produce the corrected alignment (CIGAR: 527M10S), in which the remaining 10-nucleotide soft-clipped sequence corresponds to the true non-templated 3′ tail. **(b)** Annotation of long 3′ extensions by BLAST search (section 3.9, step 2; see Note 17). When the right-clipped region exceeds 50 bp (here, CIGAR: 512M65S; 65-nt extension), it is searched by BLAST against the mitochondrial genome to distinguish biological events—such as translocation arising from *trans*-splicing or DNA recombination—from PCR artifacts such as mis-priming of ribosomal cDNA fragments. Reads with a mitochondrial BLAST hit are annotated with a translocation flag; those with no hit are searched against a cDNA database and flagged as abnormal for manual inspection.

### 3.9 Calling RNA features

The mapped consensus sequences are analyzed to identify RNA editing sites, determine 3’ end positions, and characterize 3’ terminal extensions on a per-molecule basis.

1. Call variants (**Fig. 2** step 08). LoFreq ***[28]*** is used to identify single-nucleotide variants in the mapped reads. For the *ccmC* analysis, variants are filtered for C>T changes (corresponding to C-to-U RNA editing in the transcript; **Fig. 3b**) using bcftools ***[27]***. The detected editing positions are recorded for use in the tail correction step below (*see* **Note 15**). `L3Rseq variants --input <INPUT_DIR> --outdir <OUTPUT_DIR> --ref <reference.fasta> --pattern <e.g. CT>`
2. Correct 3’ tail alignments (**Fig. 2** step 09). Each mapped read is individually processed to resolve the 3’ soft-clipped region—the sequence extending beyond the reference endpoint. The CIGAR string is parsed to identify the right-clipped portion, and a base-by-base comparison is performed between the clipped sequence and the downstream reference, tolerating mismatches at positions known to undergo RNA editing (**Fig. 4a**). The comparison proceeds until a non-editing mismatch or the end of the reference is encountered, at which point the CIGAR string is rebuilt with updated match and soft-clip counts. This correction is necessary because edited positions near the 3’ end would otherwise appear as mismatches against the genomic reference, causing premature assignment of the 3’ boundary (*see* **Note 16**). Right-clipped sequences exceeding 50 bp are additionally searched by BLAST ***[29]*** against the mitochondrial genome to detect potential trans-splicing or translocation events; sequences with no mitochondrial hit are flagged as abnormal and set aside for manual inspection (**Fig. 4b**; *also see* **Note 17**). The corrected alignments are annotated with custom SAM tags recording the 3’ end position on the reference (3E), remaining tail length and sequence after correction (RC, RS), translocation status (TL), and RNA editing counts (EC). `L3Rseq correct --input <INPUT_DIR> --outdir <OUTPUT_DIR> --ref <reference.fasta> --var < observed_variants.txt> --pattern <e.g. CT>`
3. Export per-molecule annotations to CSV (**Fig. 2** step 10). The annotated alignments are converted to a flat CSV file with one row per molecule (**Fig. 3c**). Each row contains the standard SAM fields together with the custom annotations: 3’ end position, 3’ tail length, 3’ tail sequence, translocation flag, editing count, matched alignment length, and all detected variants. This table serves as the primary output for downstream statistical analysis of the relationships among RNA processing events within individual molecules (*see* **Note 18**). `L3Rseq export --input <INPUT_DIR> --outdir <OUTPUT_DIR>`

## 4 Notes

1. PacBio sequences single strand circular DNA formed by ligating SMRTbell adapters to double strand linear DNA (< 25 kbp) ***[7]***. To maximize sequencing capacity, PacBio has developed a library preparation technique (MAS-seq ***[30]***, later renamed Kinnex ***[5, 31]***), which concatenates 8, 12 or 16 (depending on the kit) cDNA amplicons into a single long (∼ 20 kbp) fragment. The balance between the amplicon size and the concatenation factor needs to be taken into consideration. Kinnex kits can be considered for sequencing amplicons if the average size falls into the range 200 bp – 3 kbp ***[31]***.
2. Library preparation kits that support the ligation chemistry should be selected (e.g. the Ligation Sequencing kit (SQK-LSK114) and the Native Barcoding Kit (SQK-NBD114) for R10.4.1 flow cells). The Native Barcoding kit is recommended if distinction between different samples are required upon pooling or reuse of the flow cell.
3. AMPure XP beads are supplied with library preparation kits. SPRIselect beads could be used instead which enables stricter control in size selection.
4. Manuals from ONT are frequently updated, so it is highly recommended to check the latest version.
5. If bead clean-up is performed at step 1(d) in section 3.4, larger volumes of end-prepped DNA can be used as input in the next native barcoding step. This will lower the requirement for initial DNA input.
6. The scale of the native barcoding reaction may need to be increased if only a few samples are to be sequenced. Each barcoded sample contributes a fraction of the total pooled DNA; when fewer samples are pooled, the final adapter-ligated library may contain insufficient fragments. Underloading the flow cell results in low pore occupancy, which reduces both data output and flow cell longevity. For R10.4.1 flow cells, ONT recommends loading 35–50 fmol of adapter-ligated library, or approximately 100 fmol for short amplicons (< 1 kb). To compensate for low sample numbers, increase the volume of end-prepped amplicon per barcoding reaction (adjusting other reagent volumes proportionally) so that the pooled, adapter-ligated library meets the recommended loading amount.
7. Washing with SFB involves resuspending the beads, while washing with 80% ethanol does not require resuspension.
8. An NVIDIA GPU with sufficient VRAM is required for high-accuracy basecalling (HAC or SUP mode, *see* **Note 9**). Dorado basecalling uses neural network models whose computational demands scale with accuracy: the fast model can run on CPU for small datasets, but HAC and SUP models require GPU acceleration to complete in a practical timeframe. For live basecalling during sequencing, the GPU must keep pace with data acquisition; a mid-range GPU (e.g. RTX 5060 or above, with at least 8 GB VRAM) is generally sufficient for a single MinION flow cell. If live basecalling is not feasible, raw data can be saved in POD5 format and basecalled post-run.
9. MinKNOW is the operating software that controls the sequencer and can perform live basecalling during the run. EPI2ME is the companion analysis platform for interpreting ONT sequencing data. Dorado is the current ONT basecaller integrated in MinKNOW and EPI2ME. The SUP model is recommended for L3R-seq because single-nucleotide accuracy is critical for RNA editing analysis. The --min-qscore flag filters reads below a mean quality threshold; the default MinKNOW threshold for the SUP model is Q10.
10. The full pipeline can be run with a single command: L3Rseq run –input <DIR> --outdir <DIR> --ref <ref.fasta> --probe <probe.fasta> ----pattern <e.g. CT>. Use --`start-at` and `--stop-axt` to run a subset of steps. For example, `--start-at 4` skips the preprocessing steps (section 3.7 step 1; **Fig. 2** steps 01–03) for users with pre-demultiplexed data. A step-by-step example script is provided in the repository (`examples/demo_full_pipeline.sh`).
11. The default adapter sequences in `L3Rseq trim` and the RPI barcode file in `L3Rseq demux` are specific to the library preparation described in sections 3.1–3.3. Users with different adapter designs should provide their sequences via the `--adapter-fwd`, `--adapter-rev`, and `--rpi-fasta` flags. An optional pre-filtering step (`L3Rseq filter`) is also available, which rough-maps reads to the reference with minimap2 and retains only on-target reads before UMI processing. This can be invoked as a standalone command or via the `--prefilter` flag in `L3Rseq run`.
12. The UMIC-seq cluster test evaluates alignment score thresholds across a range (default: 15–29 in steps of 1) on a random subset of reads (default: 50) to identify the threshold that best separates true UMI groups from sequencing noise. The full clustering then applies this threshold to all reads. The minimum cluster size (default: 4) determines the minimum number of reads required to form a UMI group; smaller clusters are discarded as they may represent PCR artifacts or chimeric reads. Key parameters can be adjusted with `--aln-thresh`, `--size-thresh`, and `--cluster-steps`. An alternative UMI processing method based on the longread_umi pipeline [16] is available via the `--method longread-umi` flag.
13. The number of Racon polishing rounds (default: 4) can be adjusted with `--rounds`. Increasing the number of rounds may improve consensus quality for low-coverage clusters but with diminishing returns. The minimap2 preset `lr:hq` is optimized for long, high-quality reads and is appropriate for consensus polishing. Consensus quality depends on the number of raw reads per UMI cluster; clusters with fewer than 4 reads (the default `--size-thresh`) are excluded at the clustering stage to ensure sufficient coverage for error correction.
14. The target extraction step retains sequences between the forward and reverse target sequences. For *ccmC*, the defaults retain the coding sequence plus the 3’ downstream region including any non-templated extensions. The error rate (default: 0.2) and minimum overlap (default: 52 bp) control the stringency of target sequence matching. Users analyzing shorter amplicons may need to reduce the minimum overlap via `--min-overlap`. Reads that do not match both target sequences are written to a separate file (`extracted_uncut.fa`) and excluded from downstream analysis.
15. LoFreq is a variant caller designed for low-frequency variant detection in deep sequencing data. The `--pattern CT` flag filters the output for C>T variants, corresponding to C-to-U RNA editing. The `--min-af` threshold (default: 0.01) sets the minimum allele frequency for a variant to be reported. The resulting list of editing positions (`observed_variants.txt`) is used by the tail correction step to distinguish true editing sites from random mismatches. To detect other types of RNA editing (e.g. A-to-I, which appears as A-to-G in sequencing data), the `--pattern` flag can be changed accordingly.
16. In L3R-seq, the reference sequence represents the genomic (DNA) sequence. Because C-to-U RNA editing changes the transcript relative to the genome, edited positions near the 3’ end of the aligned region appear as mismatches when compared to the reference. Without correction, the aligner would soft-clip these positions, assigning a shorter aligned region and a longer 3’ tail than the true values (**Fig. 4a**). The walk correction tolerates C-to-T mismatches at known editing positions, effectively extending the aligned region through edited bases to accurately locate the true 3’ end. A user-supplied file of known editing positions (`--var`) can be used in addition to or instead of the positions detected in step 1 of section 3.9, for example when a control sample with established editing sites is available.
17. Right-clipped sequences longer than 50 bp (adjustable with `--clip-thresh`) are searched by BLAST (**Fig. 4b**) against the mitochondrial genome (default: TAIR10 ChrM) to detect reads spanning a junction between two distant genomic loci, indicative of trans-splicing or translocation. Reads with a ChrM hit are annotated with a translocation flag (`TL=1`). Reads with no ChrM hit are further searched against the TAIR10 cDNA database; those matching elsewhere are classified as abnormal and written to a separate file (`abnormal_rightclip.sam`) for manual review. Users working with organisms other than *Arabidopsis thaliana* should specify appropriate BLAST databases via the `--blast-db` and `-`-blast-db2 flags.
18. The CSV output columns include standard SAM fields (QNAME, FLAG, RNAME, POS, MAPQ, CIGAR, RNEXT, PNEXT, TLEN, SEQ, QUAL) and custom annotations: ThreePrime_end (3’ end position on the reference), ThreePrime_tail_length, ThreePrime_tail_seq, translocation (0 or 1), double_sorter (a sorting key), C-to-T (editing event count), matched_length (aligned length after correction), and All_mismatches (semicolon-separated list of all detected variants). This flat table format facilitates downstream analysis in R, Python, or spreadsheet software without requiring SAM parsing tools.

## Acknowledgements

This work was supported by Grants-in-Aid for Scientific Research (KAKENHI) from the Japan Society for the Promotion of Science (JSPS) (Grant Numbers 17J05722, 21K20658, 22H02641, 23K23904, and 24H02273).

## Notes

https://github.com/akihitomamiya-del/L3R-seq/

https://akihitomamiya-del.github.io/L3R-seq/

### Competing Interest Statement

The authors have declared no competing interest.

